# Weak interactions between groups and physical drivers of community dynamics in coastal phytoplankton

**DOI:** 10.1101/171264

**Authors:** F. Barraquand, C. Picoche, D. Maurer, L. Carassou, I. Auby

## Abstract

Phytoplanktonic communities maintain a high diversity in a seemingly homogeneous environment, competing for the same set of resources. Many theories have been proposed to explain this coexistence despite likely competition, such as contrasted responses to temporal environmental variation. However, theory has developed at a faster pace than its empirical evaluation using field data, that requires to infer biotic and abiotic drivers of community dynamics from observational time series. Here, we combine autoregressive models with a data set spanning more than 20 years of biweekly plankton counts and abiotic variables, including nutrients and physical variables. By comparing models dominated by nutrients or physical variables (hydrodynamics and climate), we first explore which abiotic factors contribute more to phytoplankton growth and decline. We find that physical drivers - such as irradiance, wind, and salinity - explain some of the variability in abundances unexplained by biotic interactions. In contrast, responses to nutrients explain less of phytoplankton variability. Concerning biotic drivers of community dynamics, multivariate autoregressive models reveal that competition between different groups (at the genus level for most) has a much weaker effect on population growth rates than competition within a group. In fact, the few biotic interactions between genera that are detected are frequently positive. Hence, our system is unlikely to be best represented as a set of competitors whose differing responses to fluctuating environments allow coexistence, as in “paradox of the plankton” models with a storage effect or a relative nonlinearity of competition. Coexistence is more likely to result from stabilizing niche differences, manifested through high intragroup density-dependence. Competition between planktonic groups and nutrients are often invoked as drivers of phytoplankton dynamics; our findings suggest instead that more attention should be given to the physical structure of the environment and natural enemies, for coastal phytoplankton at least.

## Introduction

What maintains the diversity of species-rich communities is an old yet still challenging question for ecological theory (Hutchinson 1961; Jewson et al. 2015; Li and Chesson 2016). The continued coexistence of phytoplanktonic taxa is amongst the most puzzling: species, genera and even classes of plankton are sometimes competing for the same limited resources in a seemingly homogeneous environment (Titman 1976; Tilman et al. 1982). This state of affairs led Hutchinson (1961) to refer to the “paradox of the plankton”, a paradox for which many theoretical answers have been proposed (Record et al. 2014). Classic mechanisms for biodiversity maintenance, such as niche separation or spatial variation in the environment, are not seen as obvious in the case of plankton, because species apparently compete for the same set of resources in environments that can be rather well-mixed in the absence of strong stratification of the water column (Huisman et al. 1999b).

Neutrality, i.e., per capita equivalence of birth and death rates, is another potential explanation (Hubbell 2001). There are varied life-histories in phytoplankton (Litchman and Klausmeier 2008), but neutrality does not require organisms to be equal, just that trade-offs in life history traits equalize their fitness (Hubbell 2001; Doncaster 2009). However, experimental work does suggest a difference in net reproduction rates among plankton species under varying nutrient concentrations (Tilman et al. 1982), which means that birth and death rates are in fact likely to vary a lot depending on environmental conditions.

Another leading hypothesis relates to the temporal variation in the environment (Hutchinson 1961; Chesson 2000; Litchman and Klausmeier 2001; Li and Chesson 2016). There is often a rather strong seasonal variation in the environment experienced by plankton (e.g., temperature and irradiance). A number of abiotic variables have some extra temporal variation as well, often due to perturbations from nearby ecosystems, especially in coastal systems like estuaries and lagoons (e.g., nitrogen (N), phosphorus (P), silicon (Si), due to terrestrial inflows). Theory posits that diversity maintenance can be precisely due to this temporal variation in the environment (Chesson 2000; Li and Chesson 2016), as in models for the storage effect (Chesson and Huntly 1997). Such models are different from neutral models, as species have differentiated responses to environmental variables. Some near-neutral models do include temporal responses to the environment (Kalyuzhny et al. 2015), but near-neutral models usually assume that species compete in a zero-sum game, where no species can increase without a decrease in some other species (but see Jabot and Lohier 2016). However, total biomass across species/genera often fluctuates across several orders of magnitude throughout the year, blooms being the most obvious defining feature of planktonic dynamics. This makes phytoplanktonic communities fundamentally different from other species-rich communities, such as forests or coral reefs, where the fluctuations in biomass are clearly milder and near-neutral dynamics may be more plausible (Hubbell 2001; Segura et al. 2017; we will come back to the likelihood of neutral dynamics for phytoplankton in our discussion).

Temporal variation is therefore a likely culprit for coexistence in plankton - as initially proposed by Hutchinson (1961). In this scenario, all species are assumed to outcompete others at some point in time, with alternance in the ranking of competitors preventing competitive exclusion (Descamps-Julien and Gonzalez 2005). This mechanism is likely to work if species have differing nonlinear responses to the environment (i.e., relative nonlinearity of competition), or a covariance between environmental conditions and competitive strength, as in the storage effect (Chesson 2000; Fox 2013; Li and Chesson 2016). The plausibility of a fluctuation-driven coexistence mechanism has been empirically verified on diatoms that belong to different genera (e.g., Descamps-Julien and Gonzalez 2005) though some experiments and models also suggest that planktonic communities might be in a chaotic or seasonally-driven chaotic state (Huisman and Weissing 2001; Benincà et al. 2008; Dakos et al. 2009). Such chaotic state invokes other fluctuation-driven coexistence mechanisms, with nonlinearities in functional forms promoting endogeneously-driven fluctuations. Of course, these mechanisms are not exclusive, and coexistence is generally due to the joint influence of equalizing factors generated by differential responses to the environment (i.e., species have the same fitness when averaged over the variation in the environment) and stabilizing mechanisms (i.e., species increases when rare) that make coexistence all the more likely (Chesson 2000).

In summary, diversity maintenance in planktonic communities seems essentially tied to abundance fluctuations throughout the year, due to both abiotic and biotic factors. Therefore, any framework examining competition among planktonic species should account for both the broad variations in environmental conditions and in planktonic abundances over time. Multivariate autoregressive (MAR) modeling is one such dynamic framework that has been increasingly used to examine interactions between planktonic groups in a dynamic environment (Klug et al. 2000; Ives et al. 2003; Hampton and Schindler 2006; Huber and Gaedke 2006; Scheef et al. 2013; Griffiths et al. 2015; Gsell et al. 2016). MAR models enable the estimation of interaction strengths between taxa (Ives et al. 2003; Mutshinda et al. 2009) as well as the dependence of population growth rates on abiotic variables (Hampton et al. 2013), both necessary to model planktonic dynamics. These models are linear on a log-scale, hence they represent population growth processes as multiplicative, power-law functions of densities that can approximate more complex nonlinear functional forms (Ives et al. 2003). One can also consider an increasing degree of nonlinearity using phase-based MAR models (also called “Threshold AR” models, Stenseth et al. 2015), in order to examine in detail the plausibility of nonlinear coexistence mechanisms.

Most plankton-based MAR analyses so far (but see Huber and Gaedke 2006) have aggregated data at the class level, therefore preventing any attempt to examine the strengths of competitive interactions among species or genera. Recent papers highlight that a finer taxonomic resolution would allow for a better understanding of community dynamics (Griffiths et al. 2015; D'Alelio et al. 2016). Here, we take take advantage of a long-term dataset of coastal phytoplankton monitoring (>20 years of biweekly counts) with fine taxonomic resolution - to the genus level - to investigate:

1. What are the abiotic drivers of population growth (i.e., factors that initiate or terminate blooms)? Are nutrients or hydrodynamics factors most influential?
2. What are the strengths of interactions between groups and how does variation in biotic factors affect the joint dynamics of the community?
3. What are the implications of our results, obtained on a rich diatom and dinoflagellate assemblage, for biodiversity maintenance (i.e., what coexistence mechanisms can be expected)?

## Material and methods

### Study area and sampling details

Arcachon Bay (AB) is a 155 km^2^ coastal lagoon located in the south-west of France (Fig. 1). AB is connected to the Atlantic Ocean by two main channels (41 km^2^), and total freshwater inflow distributes between the Eyre river (83%) and Porge canal (11%), rainfall and ground-water (6%, Rimmelin et al. 1998). Using a 2D-hydrodynamics model, Plus et al. (2009) estimated AB's tidal prism, i.e. the volume of water leaving the bay at ebb tide, around 384 million m^3^, which can be compared to the much lower maximum daily water discharge of 12 million m^3^ from the Eyre, the main river entering the bay. Plus et al. (2009) also computed a flushing time for the lagoon between 13 and 30 days and highlighted a high return flow factor (fraction between 0.94 and 0.95, 1 being the maximum value), meaning that a compound carried out of the lagoon is likely to return to it. The environment is therefore well-mixed at large spatial scales, and mostly marine-influenced. In addition, the wind regime has a strong influence on the hydrodynamics of the bay: westerly winds represent about 77% of total wind and can increase water residence time in the bay up to 10 days, leading to surges up to 50 cm (Rimmelin et al. 1998; Plus et al. 2009, 2015). The oceanic temperate climate is characterized by an amount of precipitation averaging 785 mm year^−1^, a total irradiance around 475 kJ cm^−2^ year^−1^ and an average water temperature around 15.6°C, with a strong seasonality for all parameters (rainfall are 40% less abundant in summer than in winter, irradiance is 4-time stronger and temperature varies between 9°C and 23°C). More details on AB (e.g., absence of stratification of the water column) are given in Appendix S1: Section S1.1.

**Figure 1:**
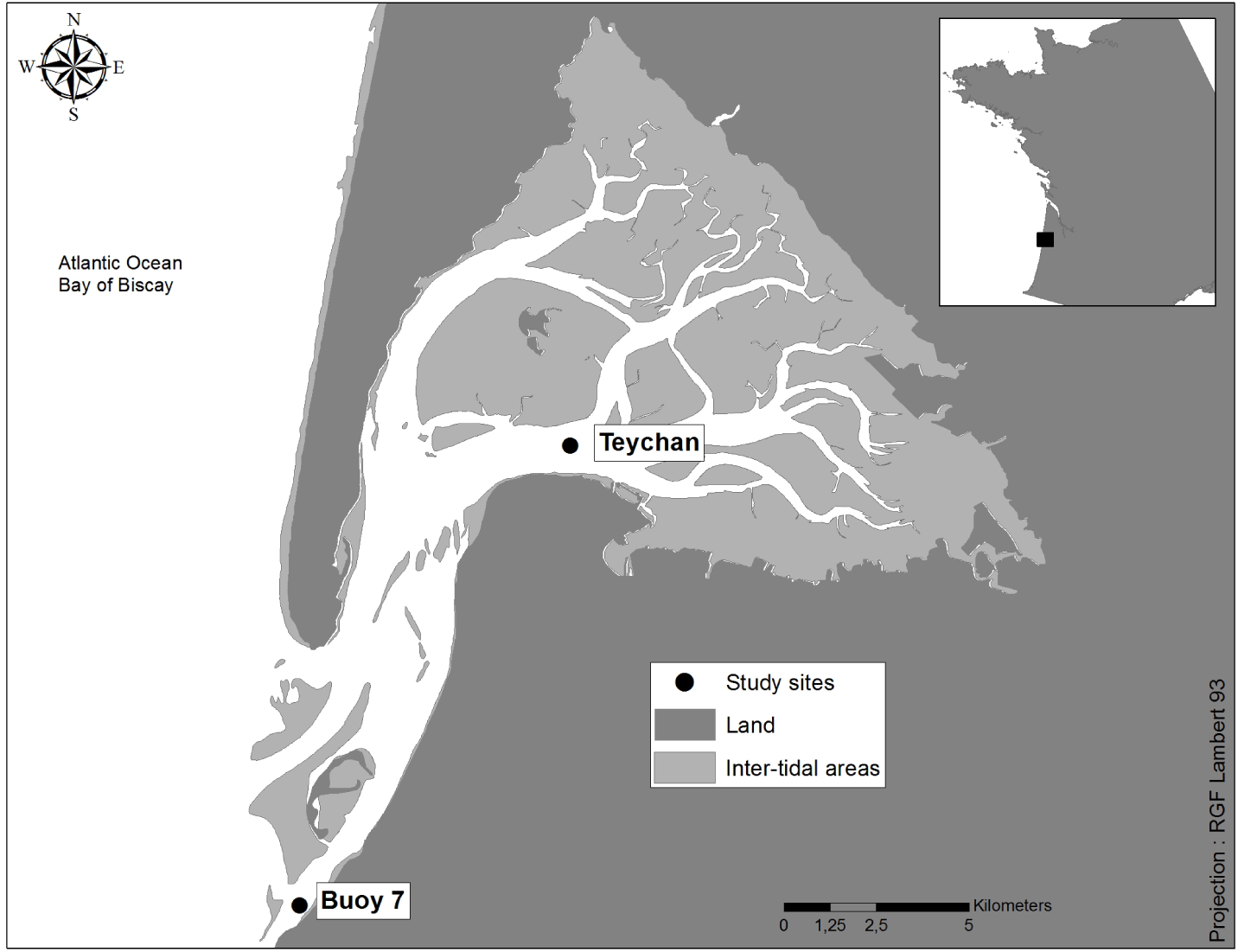
General view of Arcachon Bay and location of sampling points (Teychan and Buoy 7) within the study area. Exact coordinates and further information about the sampling sites are given in the main text.

Water samples were collected at 2 sites: Teychan (44°40'25” N / 1°09'31” W, water depth = 17.30m) and Buoy 7 (B7, 44°32'32” N / 1°15'49” W, water depth = 14.5m), from 1987 to 2015 (709 dates) and 2003 to 2015 (311 dates), respectively. For more information on water sampling, see Appendix S1: Section S1.2. Teychan is located near the limit drawn by Bouchet (1993) between water masses influenced by oceanic waters or by continental inputs, in the middle of AB (Fig. 1), while Buoy 7 is at the oceanic entrance of the channel. Teychan is therefore expected to be more sensitive to continental inputs while Buoy 7 should be mainly subject to marine influences (Bouchet 1993; Glé et al. 2007).

### Plankton ecology in Arcachon Bay

There is a wide temporal variability in AB, with an early phytoplankton bloom in spring and usually a late one in autumn. Based on a five-year spatialized field study of micro-phytoplankton primary production, Glé et al. (2007) concluded that part of the marine plankton in AB may be coming from the adjacent Bay of Biscay oceanic waters, and the bloom within the lagoon could therefore be “seeded” by the ocean. According to their study, marine plankton find favorable conditions in AB as nutrient stocks from freshwater inflows are higher than in the ocean. A low turbidity, associated with a well-mixed water column (see Appendix S1: Fig S2), improves light penetration and favors a rapid population growth in response to elevated irradiance happening as early as the end of February, depending on climatic variability. According to Glé et al. (2008), increases in plankton abundance lead to nutrient depletion that is reflected in a lower productivity during summer. Both experimental (Glé 2007) and modeling (Plus et al. 2015) studies have shown so far that N might be limiting during summer in the interior of AB, while P restriction might be the main driver of productivity in external waters (Glé et al. 2008). On a shorter timescale, daily ebb and flow participate in nutrient cycling (amounting to 55% P and 15% N in the bay according to Deborde et al. 2008) while annual plankton loss to the ocean is estimated between 383 and 857 tC year^−1^ by Plus et al. (2015). A switch from early, large plankton assemblage with low biodiversity to a more diverse and smaller-bodied community has been noticed by Glé et al. (2008) although the latter study was restricted to a single year (2003), characterized by a dry winter and a strong summer heat wave. AB is also characterized by a higher average salinity and lower turbidity than other French Atlantic coastal bays where biotic and abiotic factors are similarly monitored (David et al. 2012).

### Phytoplankton data

The National Phytoplankton and Phycotoxin Monitoring Network (REPHY^1^) involves plankton collection every two weeks, within two hours at high tide (for more details, see Appendix S1: Section S1.2). Data collection started in 1987 in AB (Teychan), so the full dataset consists in 709 sampling dates. The full dataset is used in all analyses except for multivariate autoregressive models including all planktonic groups (see “MAR(1) models”): as cryptophytes were not recorded properly before 1996, and this could affect the counts of other taxa, we focused on observations post-1996 for MAR(1) models.

Taxonomic units were aggregated at the genus level following previous work on this and other plankton datasets (Hernandez et al. 2013; Hernández Fariñas et al. 2015) according to the plankton experts’ knowledge (see Acknowledgements). We made only two exceptions to this “genus rule”: cryptophytes and euglenophytes could not be consistently identified below the class level, but their abundance throughout the time series justified to keep them in the analyses.

Our analyses therefore focused on the best-resolved and abundant planktonic groups over time (Table 1). Hereafter, we remained consistent with the terminology used in population dynamics theory and used the word “population” for groups described in Table 1. The term “community” encompassed all planktonic taxa interacting in AB.

**Table 1:** Name and composition of the planktonic groups used in the paper, with their average proportion in the planktonic community, calculated as the ratio of their summed abundance over the summed abundance of all identified planktonic organisms, and the frequency of detection, calculated as the ratio of samples in which the group was present over the total number of samples over the whole time series at Teychan and Buoy 7 in Arcachon Bay

| Code | Groups | Average fraction<br>in the community<br>(%) | Frequency of<br>detection<br>(%) |
| --- | --- | --- | --- |
| AST | *Asterionella* + *Asterionellopsis* +<br>*Asteroplanus* | 0.19 | 0.80 |
| CHA | *Chaetoceros* | 0.12 | 0.89 |
| CRY | *Cryptophytes* | 0.44 | 0.64 |
| EUG | *Euglenophytes* | 0.01 | 0.64 |
| GUI | *Guinardia* | 0.03 | 0.73 |
| GYM | *Amphidinium* + *Gymnodiniaceae* +<br>*Gyrodinium* + *Katodinium* | 0.01 | 0.73 |
| LEP | *Leptocylindrus* | 0.12 | 0.67 |
| NIT | *Ceratoneis* + *Nitzschia* +<br>*Hantzschia* + *Bacillaria* | 0.03 | 0.94 |
| PRP | *Protoperidinium* + *Peridinium* | 0.01 | 0.78 |
| PSE | *Pseudo-nitzschia* | 0.06 | 0.66 |
| RHI | *Neocalyptrella* + *Rhizosolenia* | 0.01 | 0.63 |
| SKE | *Skeletonema* | 0.06 | 0.62 |

Plankton abundance values (cell L^−1^) were linearly interpolated over a regular time sequence with a 14-day time step to correct for some sampling irregularity (following Hampton et al. 2006). Missing values observed during blooms and spanning 2 time steps or less were replaced by an interpolated value, as these missing values were almost surely not zeroes, but a lack of detection. Remaining dates for which no individual was observed for a given group were treated as missing values instead of indicators of null abundance. We evaluated several methods for reconstructing time series without gaps and they are detailed in Appendix S1: Section S2.1; we followed Hampton and Schindler (2006).

### Abiotic data

Environmental variables collected at Teychan and Buoy 7 included water temperature (TEMP in °C), salinity (SAL, in g kg^−1^), nutrients (ammonia NH4^+^, silicates Si(OH)_4_, phosphates 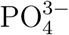 and nitrates-nitrites NO_x_=NO_2_ + NO^3−^, all in µmol L^−1^) and suspended particulate matter (SPM, in mg L^−1^), with an organic part assessment (SPOM, in mg L^−1^). They were measured following Aminot and Kérouel (2004, 2007) (see Appendix S1: Section S1.2 for more details).

Some abiotic variables (nutrients and SPM) could not be recorded at Teychan until 2007. Their values were therefore replaced by measurements at the closest hydrological station, Tès (44° 39'59 N / 1° 08'40 W; 1.4 km from Teychan), which shows a similar hydrodynamic functioning (see Appendix S1: Fig. S3). For Buoy 7, all abiotic variables could be measured onsite for each year.

Daily meteorological data -rainfall (mm), irradiance (J cm^−2^), wind direction and velocity (m s^−1^) - were provided by Météo France for the nearby Cap Ferret station (44° 37’ N / 1° 14’ W) and were used as inputs for both sampling stations. Wind energy was extracted from these data as the squared velocity (m^2^ s^−2^).

Daily North Atlantic Oscillation (NAO) index was downloaded from the National Oceanic and Atmospheric Administration (NOAA) Weather Service for Climate Prediction Centre website^2^. Monthly Atlantic Multidecadal Oscillation (AMO) index was downloaded from the NOAA Physical Science Division website^3^.

All environmental data were linearly interpolated on the same dates as plankton sampling dates, removing missing values (see Appendix S1: Fig. S3).

We also transformed abiotic variables to represent meaningful biological and physical processes. We summed 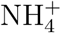 and NO_x_ as nitrogen suppliers (Ntot). We integrated inflow (CumDebit), rainfall (CumPrec) and irradiance (CumRg) between *t −* 1 and *t* to represent the growth conditions of plankton groups between two sampling dates (Glé et al. 2007). NAO was averaged over the same period and a variable summarizing the effect of wind was constructed using the mean wind energy over 3 days (F. Ganthy, Ifremer, personal communication).

All datasets and R scripts for analyses are available online in a GitHub repository^4^.

### Environmental drivers of population dynamics

The high number of possible explanatory variables with some degree of auto-correlation and cross-correlation (see Appendix S1, Figs. S4 - S7) led to a collinearity problem indicated by a very high condition index (CI=150, well above the thresholds of 10 or 30 indicated by Belsley 1991). To select relevant variables, our approach was three-fold: (a) spectral analyses enabled us to detect significant coherency (i.e., correlation in the spectral domain) between biotic and abiotic cycles to select variables that seemed coupled to plankton abundance, (b) nutrients being highly correlated (Appendix S1: Fig. S4-Fig. S7), we selected the most relevant ones based on literature values on planktonic requirements, which then led us to (c) compare a nutrient-limitation model using N and P as covariates (including the possibility for saturating functions of nutrients) to a physics-driven model using irradiance, salinity and wind energy as covariates. This comparison was based on log-linear, single species autoregressive modeling. A “full model” including both types of drivers was also compared to the aforementioned more parcimonious models. Climate indices (NAO and AMO) formed a third group of variables for which we developed a monthly model. Coefficients associated to climate indices were low and explained little the observed variation, which was in line with previous results for AB (David et al. 2012); we therefore chose to discard such large-scale indices from further analyses, as they were also more difficult to interpret ecologically.

### Spectral analyses

Plankton groups show periodic cycles that can be related to seasonal cycles. Correlating short- and long-term cycles for both variable types can help extract planktonic responses to environmental variability from process noises. For this, we compared each genus periodogram, using a modified Daniell kernel with a smoothing window of 2 time steps (1 month) and no tapering, with abiotic variables. Coherence significance was determined with a corrected (Bonferroni) 5% threshold.

### Nutrient limitations

Nutrient uptake is a highly variable process both within and between plankton genera (Paasche 1973; Fisher et al. 1988; Sarthou et al. 2005; Reynolds 2006; Litchman et al. 2007). To assess potential limitations to growth and insert ecologically meaningful predictors into autoregressive models, we used recently published half-saturation values for both marine diatoms and dinoflagellates found in Litchman et al. (2007). These were used to create saturating functions of nutrient concentrations. Concentrations were considered saturating when twice superior to 0.65 µM L^−1^, 1.25 µM L^−1^ and 1.0 µM L^−1^ for P, N and Si, respectively, for diatoms, and 1.4 µM L^−1^ and 7.0 µM L^−1^ for P and N, respectively, for dinoflagellates. We averaged values for cryptophytes from two models (Hood et al. 2006; Rigosi et al. 2011): saturating values were twice above 1.2 µM L^−1^ and 2.7 µM L^−1^ for N and P, respectively. Based on available literature, nutrient requirements for the euglenophytes could not be so thoroughly assessed: average half-saturation values from Fisher et al. (1988) were used for N (2.0 µM L^−1^) and from Chisholm and Stross (1976) for P (1.4 µM L^−1^).

### Single-species autoregressive models

Before using multivariate autoregressive (MAR) models, each planktonic group was studied on its own. The modeling philosophy adopted here classically assumes that a multidimensional dynamical system might be represented as a delayed unidimensional dynamical system (Takens 1981; Turchin 2003), even in a stochastic context (Abbott et al. 2009). Accordingly, each population can be described by an autoregressive (AR) model where the number of time lags depends on the dimensionality of the system. The goal was to identify potential abiotic drivers of population dynamics before the multivariate analyses, in order to fit later on more parsimonious MAR models (Fig. 2; bear in mind that a full-matrix MAR(1) model with covariates can require to estimate up to 204 parameters).

**Figure 2:**
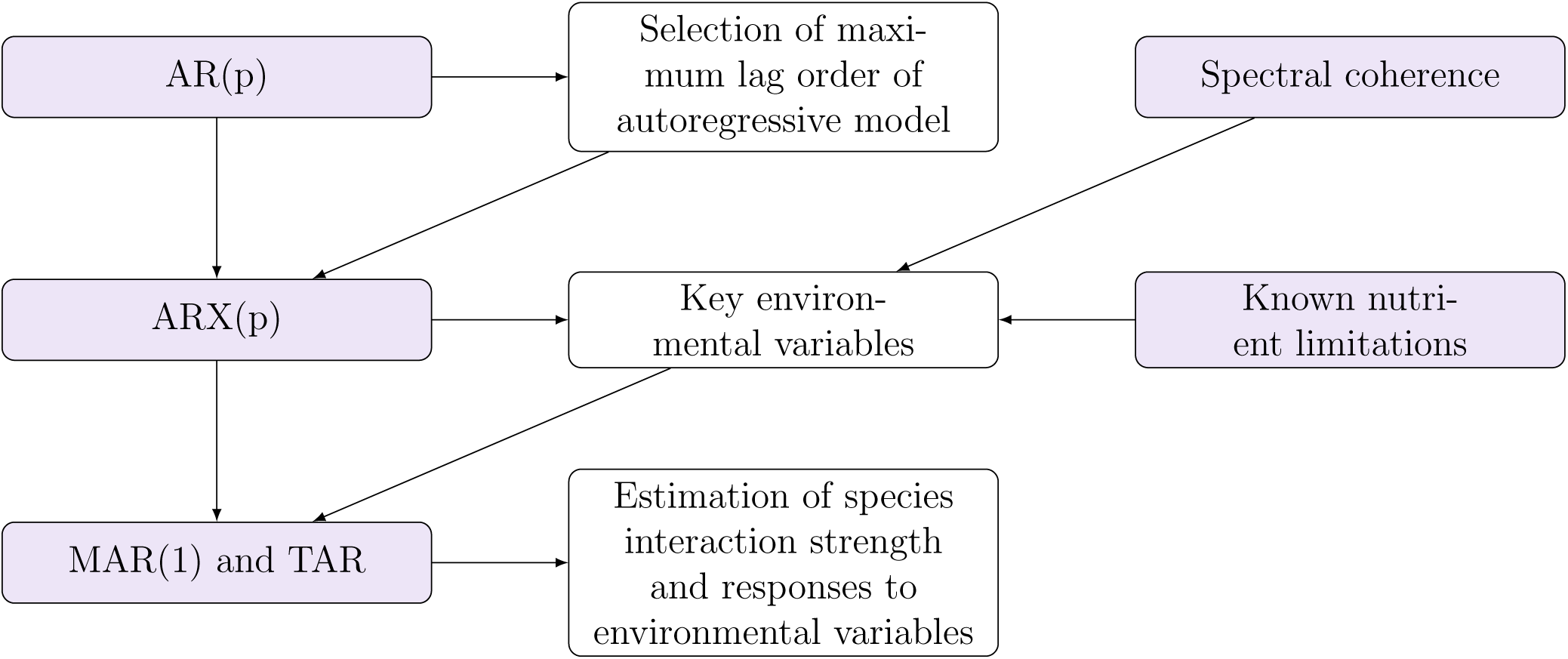
Workflow of the main statistical analyses.

#### Lag order selection

For log-linear AR modeling, focusing on one species at a time, we assumed that the plankton's ecology did not vary greatly between sites and concatenated data from both stations to increase the effective sample size in the statistical analyses. We fitted a log-linear AR(*p*) model, which allowed the determination of the maximum lag order *p*. The shape of growth-abundances curves (see Appendix S2: Fig. S1) and analyses of residuals (Appendix S2: Section S2.5) confirmed that log-linearity on a log-scale - Gompertz rather than Ricker growth (Ives et al. 2003) - was an appropriate approximation. We used the `arima` function from the `stats` package (Venables and Smith 2013) in R (version 3.3.2) to estimate coefficients and corresponding Akaike Information Criterion (AIC) for each AR(*p*) model. The number of time lags considered varied between 1 and 30, the maximum reasonable value for the length of our dataset (780 dates for taxa monitored after 1996). The likelihood was estimated using the Broyden-Fletcher-Goldfarb-Shanno (BFGS) optimization method, an approximation of Newton's method, with a maximum of 100 iterations.

#### Adding environmental variables

AR(p) models with abiotic predictors affecting growth rates - sometimes called ARX(p) - were then used to study the environmental effects on planktonic growth, according to equation 1.

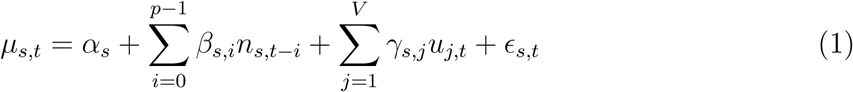

where *µ_s,t_* = *n*_*s,t*+1_ *− n_s,t_* is the population growth rate on a log scale for genus *s*, *α_s_* is the genus-specific productivity when population size is zero on a log scale, *β_s,i_* is an AR coefficient that can represent intragroup density-dependence at lag *i*, *u_j,t_* is environmental variable *j* for time *t*, with effect *γ_s,j_* on the population growth rate *ε_s,t_* is a random variable modeling both environmental and observation error with mean 0 and variance *σ*^2^ (Dennis et al. 2006).

The maximum number of time lags was homogenized between species: our results showed that for AR(p), a 3^*rd*^ order AR model minimized ∆AICc = AICc − AICc_min_ for most species and resulted in 4 models out of 12 falling in the range [AICc_min_, AICc_min_ + 2] and 7 more within [AICc_min_, AICc_min_ + 4]. Therefore, *p* = 3 was enough to capture the dynamics using single-species models. Models with *p* = 1 resulted in similar coefficients but slightly higher AICc values.

All abiotic variables were standardized in order to make their effects comparable. As Si and N were highly correlated (Appendix S1: Fig. S5, S6; a strong lagged-correlation also appeared in Fig. S7(d)) and Si seemed less limiting than N (Table 2), P and N were the only nutrients retained in linear ARX(3) models. Two models were used to assess N availability: the first one used raw concentrations while the other one used a saturation function (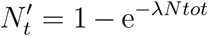, with in accordance to N half-saturation constant) to lower the impact of an increase in concentrations above a certain threshold. We also considered models using nutrient ratios (Si/N and P/N) instead of raw concentrations. All abiotic variables were then transformed to take into account seasonality. First, a seasonal component was computed from a linear regression of temperature data against trigonometric functions with an annual frequency (Appendix S1: Section S1.7). Abiotic variables were then regressed against this seasonal component and residuals from this regression were added to the model as abiotic variables (see Eq. 2).

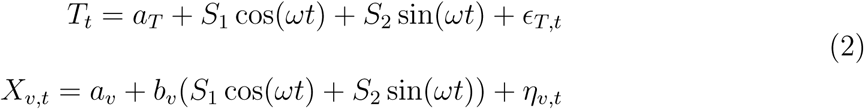

where *T_t_* is the temperature at time t, *S*_1_ cos(*ωt*) + *S*_2_ sin(*ωt*) is the seasonal component with a normal noise *ε_T,t_* and *η_v,t_* is the residual component of covariable *X_v,t_* when season has been taken into account, *a_v_* and *b_v_* being associated regression coefficients.

**Table 2:** Proportion of observations (%) at Teychan and Buoy 7 for which nutrient concentrations were above twice the half-saturation parameters found in the literature for different planktonic phyla (see the text for values and references). Dino. = Dinoflagellates.

|  | Phosphorus |  |  |  | Nitrogen |  |  |  | Silicon |
| --- | --- | --- | --- | --- | --- | --- | --- | --- | --- |
|  | Diatoms | Dino. | Cryptophytes | Euglenophytes | Diatoms | Dino. | Cryptophytes | Euglenophytes | Diatoms |
| Teychan | 0 | 0 | 0 | 0 | 60 | 17 | 61 | 50 | 96 |
| Buoy 7 | 0 | 0 | 0 | 0 | 37 | 0.6 | 38 | 26 | 57 |

Model evaluation was performed in two steps. We first used physics-based variables (salinity, irradiance and wind energy) and nutrients (nitrogen and phosphorus) together to define a “full” model including all covariables. Three representations of nutrient variables were considered and compared: raw concentrations, saturating functions of concentrations see above “Nutrient limitations” - or nutrient concentrations ratios. We considered, for each model, versions with and without a dedicated seasonal component. This led to six different full-model formulations. The minimum AICc for a majority of plankton groups determined the best full model representation. The full model was then compared to its “nutrients-only” and “physics-only” counterparts (i.e., models with only nutrient or physical variables as predictors, respectively), in order to choose the minimum amount of abiotic predictors for the following MAR(1) analysis.

### MAR(1) models

Multivariate autoregressive (MAR) models are multivariate extensions of AR models. Instead of focusing on one planktonic group at a time, these models describe the change in abundance of all groups at the same time, conditioned by past values of both the *V* environmental variables and the abundance of the *S* other planktonic groups. We used Ives et al. (2003) formulation (Eq. 3), which explains growth between times *t* and *t* + 1 by the abiotic variables at time *t*+1 (because of the rapid growth of plankton, using variables both at *t* and *t* + 1 could be valid choices, see Hampton et al. (2013) for an alternative). This formulation was chosen because it (1) led to smaller AICc and Bayesian Information Criterion (BIC) for otherwise identical models and (2) resulted in better consistency between estimates at Teychan and Buoy 7 sites (77% of covariate effects are qualitatively similar). It should be noted that such choice does not impact the qualitative results in the interaction matrix (for more discussions on the difference between Ives et al. (2003) and Hampton et al. (2013) formulations, see Appendix S3). The MAR(1) model is described by Eq. (3)

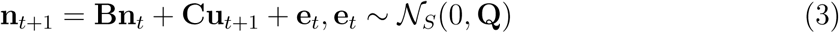

where **n**_*t*_ is a *S ×* 1 log abundance vector of planktonic genera at time t, **B** is a *S × S* community (interaction) matrix. When using such MAR(1) model, we need to substract one to autoregressive, intragroup coefficients in order to make them comparable to intergroup interaction values: no intragroup regulation leads to *b_ii_* = 1 whereas a strong intragroup effect is characterized by *b_ii_* close to or below zero (see Ives et al. (2003) for details). For *i* ≠ *j*, *b_ij_* describes the effect of genus *j* on genus *i*. **C** is a *S × V* environment matrix describing the effects of variables **u** on planktonic growth. Therefore *c_ij_* is the effect of variable *j* on genus *i*. e_t_ is a noise vector which covers both process and observation error (Appendix S1: Section S2.1, describes the handling of missing or incomplete observations), following a multivariate normal distribution with a variance-covariance matrix **Q**. In the following, **Q** was chosen diagonal. Indeed, the absence of constraints on the variance-covariance matrix could lead to 66 additional parameters, in a system that already requires to estimate between 72 and 204 parameters. We checked what changes were induced by non-diagonal **Q**, and those are minimal (see Appendix S4), thus a diagonal **Q** is clearly the parsimonious choice here.

We used the MARSS package (Holmes et al. 2012, 2013) to fit MAR(1) models. Before fitting the MAR models, we had to handle missing values in plankton data. The MAR models already had a high number of parameters to estimate, so we chose not to add more through the estimation of missing values. They were replaced by a random value between zero and the minimum value of the corresponding time series, for values that could not be interpolated, following Hampton and Schindler (2006) (see Appendix S1: Section S2.1 for alternatives). MARSS models cannot deal with missing values in the covariates: all abiotic variables thus had to be linearly interpolated (which posed no problem since abiotic variables were sampled at higher frequencies than plankton). All time series were centered and standardized. We used a physics-only model with a dedicated seasonal component to represent plankton dynamics, based on previous single-species analyses.

Five interaction matrices were considered. Except for the null model which allowed no net interactions between groups, all interaction matrices included the only documented effect in the studied planktonic community: a possible predation between dinoflagellates and cryptophytes (Moeller et al. 2016). Because all other interactions were unknown, we considered several contrasted scenarios for model fitting. Eq. (4) defines for example a scenario differentiating between pennate and centric diatoms. Three other interaction matrices are presented in Appendix S2: Section S2.3.

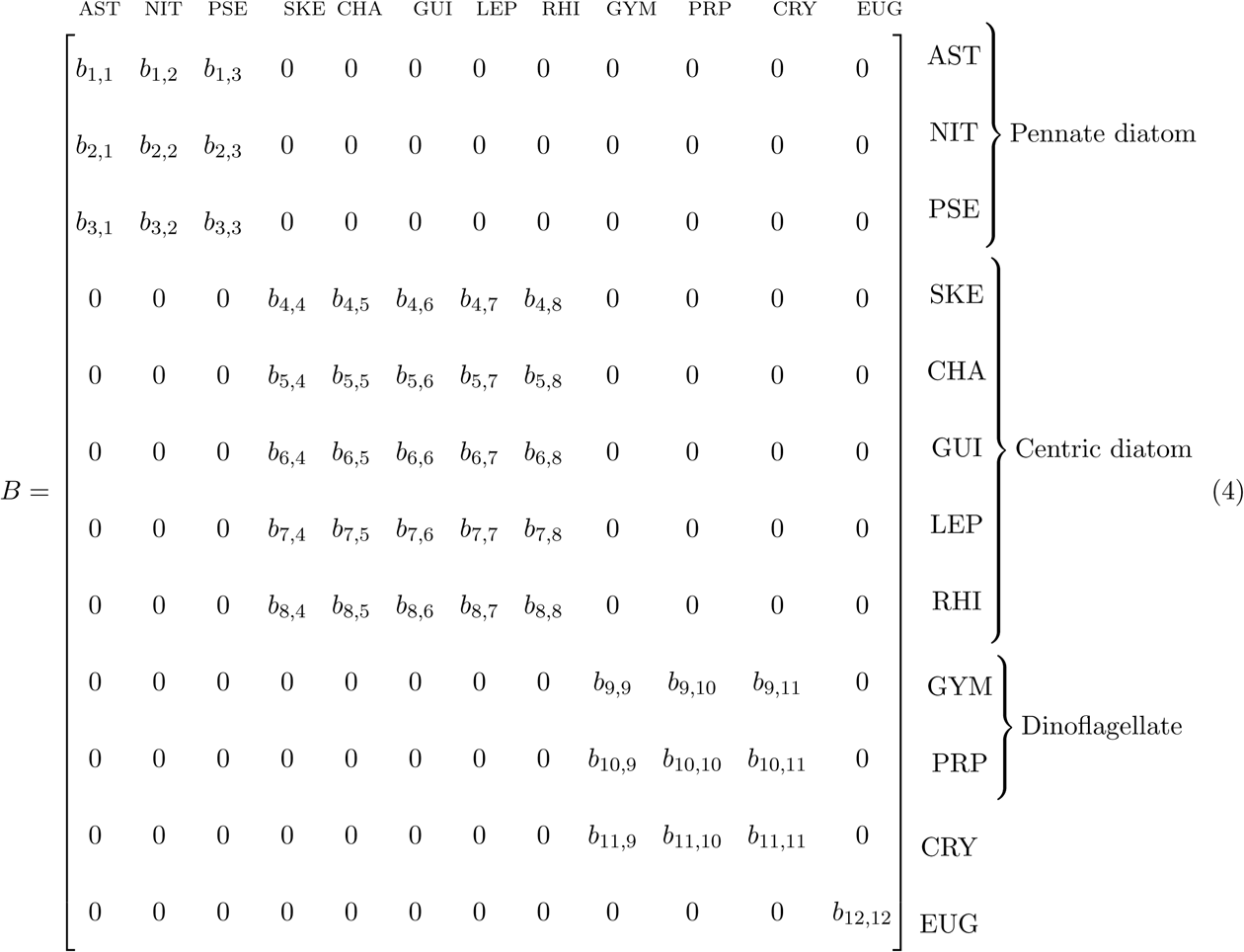

In our first scenario, “the full model”, the interactions were totally unconstrained. The second and third scenarios only took into account intra-phylum competition or facilitation (i.e., within diatoms or dinoflagellates). The third scenario added a further difference between pennate and centric diatoms (eq. 4). Finally, the fourth model only took into account inter-phylum interactions, which ecologically represents a scenario where, within a phylum, organisms are too close to each other to compete. For each scenario and model fit, 95% confidence intervals (CIs) were calculated for all parameters by boostrapping and only CIs that did not cross zero were considered significant. AICc was used to select models, and this model selection was compared to results obtained by BIC and cross-validation. Cross-validation was performed on Teychan site by applying each model to the first 10 years (1987-1996) of the Teychan time series, removing the effect of cryptophytes and euglenophytes which were not counted until 1996. This was not the case for Buoy 7 for which we used the whole time series (starting in 2003) in the estimation process.

We chose not to construct models including both observation and process error, because the magnitude and variability of observation errors were unknown, and it has been showed clearly under such conditions that introducing an unconstrained observation process in a state-space model can result in a loss of identifiability of parameters (Knape 2008; Auger-Méthé et al. 2016). Appendix S1: Section S2.1 details different types of observation errors and their handling. Parameters were estimated by default *via* maximum-likelihood using an expectation-maximisation algorithm (see Appendix S2: Section S2.2 for more details on MARSS tuning parameters used in our estimations). The optimization process relied on two convergence tests: the absolute change in log likelihood from one iteration to another and its slope *vs.* the iteration number had to be below 0.001 to accept convergence. We thoroughly tested our model selection approach using 10 simulated datasets resembling our empirical dataset, for 3 contrasted ecological scenarios: (1) effects of environment only, (2) effects of interactions only, and (3) both environment and interactions (see Appendix S2: Section S2.1). The results on simulated data showed that AICc selected the appropriate model in each scenario, thus there was no need for more complex model selection using AICb or refined procedures as did Ives et al. (2003) (results are provided in Appendix S2: Section S2.1).

### Non-linear dynamics: bivariate phase-dependent (TAR) models

Complex nonlinear dynamics (sensu May 1974; Turchin 2003) may not be completely captured by linear models, even in the logarithmic scale (Stenseth et al. 1998a, 2004). The log-linear MAR(1) framework indeed essentially assumes power-law functional forms. Modeling accurately differential responses to environmental variations like the storage effect (Chesson and Huntly 1997) or relative nonlinearity of functional forms warrant models with a finer temporal structure. Fortunately, threshold autoregressive (TAR) models (Stenseth et al. 1998b, 2004) can account for different temporal regimes by introducing different phases separated by threshold values - they are piecewise log-linear. TAR models have been successfully applied to diverse ecological studies (Stenseth et al. 1998a,b, 2004). They can be described by Eq. 5, where in our case two phases are defined:

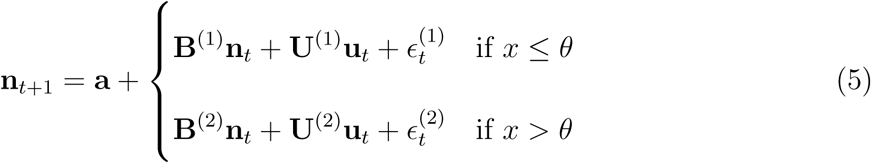

where **n**_*t*_ is the log abundance at time *t*. The parameters **B**^(1/2)^, **U**^(1/2)^ and 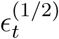 have the same meaning as in the usual MAR model but can have different values in different phases. The phase is defined by *x* which can be both an intrinsic state variable (population abundance or dynamics in Stenseth et al. 1998a; Barraquand et al. 2014) or an extrinsic variable (Stenseth et al. 2015) which is compared to a threshold *θ*. We describe below how the definition of phases and thresholds map to ecological mechanisms like the storage effect and relative nonlinearity of competition.

In order to keep the analyses tractable and the number of parameters reasonable, given the length of the time series, we focused on the two main genera in Arcachon Bay: AST and CHA. According to our findings using log-linear MAR models (see above), they are not supposed to interact, being pennate and centric diatoms respectively. We tested their interactions using the phase-based models to make sure that the absence of reported interaction was not due to hidden nonlinearities.

The first phase-based models, with phase defined through population densities (*x* is defined by AST and CHA abundance in Eq. 5), model possibly complex functional forms of the interactions between AST and CHA. Competition and response to environmental variables have then separate coefficients for high and low densities of the two plankton groups. This corresponds well to “relative nonlinearity of competition” (see e.g. Fox 2013 for model equations). The second, environment-based phase models (*x* and *θ* are defined by clusters of environmental conditions in Eq. 5) allow for coefficients to differ depending on environmental conditions. The latter models are particularly relevant to see if a so-called “storage effect” could happen, in which the effect of abiotic variables on population growth covary with the values of the environmental variables themselves (Chesson 2000). For alternative, non-parametric ways to search for storage effects, see Ellner et al. 2016.

#### Density-based phases - nonlinear competition

We defined cumulated irradiance as the main environmental driver, based on MAR(1) model results. Litchman et al. (2004) also showed that light conditions could impact competition for resources between plankton groups (results when using temperature as the main environmental effect can be found in Appendix S5: Section S1.2). Two log-linear MAR(1) models were fitted to the data, on both sides of the threshold: the first one corresponded to non-bloom conditions while the second one represented the dynamics during a bloom. The log-abundance threshold for regime switch was set to 11, which implies that blooming conditions represented about 25% of both AST and CHA time series.

#### Environment-based phases - storage effect

Defining a environmental-based TAR model when the environmental conditions are multidimensional requires two main choices: the choice of a unifying variable that can summarize external conditions of the system (i.e., the phase) and the definition or estimation of a threshold for such a variable. The number of different phases, and therefore thresholds, is itself debatable. We chose to use 2-phase hierarchical clustering based on Ward's method (see Appendix S5: Section S1.1 for details). Two phases allowed to differentiate between low chlorophyll and salinity and high chlorophyll and salinity conditions, while making results comparable to both previous sections and other works in the literature (D'Alelio et al. 2015). Using this clustering method to define a phase for both Teychan and Buoy 7 datasets, a different MAR model was fitted to each phase. As before, AICc was then used to compare between diagonal and full interaction matrices.

## Results

### Environmental drivers of population dynamics

Here, we describe the correlation patterns of environmental drivers and plankton groups in the frequency domain, that helped us, together with ecological knowledge, to select the key environmental drivers used later in ARX(p) and MAR(1) models (Fig. 2).

### Spectral correlation analyses

Salinity was related to both freshwater inflow and precipitation, and having all three variables in our final models would not be desirable given their high correlation. As salinity cycles were more often correlated with plankton dynamics (Appendix S1: Fig. S18), and this variable also makes biological sense for the plankton niche (see Discussion), we retained only salinity. Irradiance and temperature were also correlated, which was of course logical given a higher irradiance locally heats up the water temperature. Both could have been considered, but we chose to use irradiance as the leading variable summarizing solar energy. Furthermore, on purely statistical grounds, CRY, the only group with differential responses to irradiance and temperature, were more sensitive to irradiance cycles (Appendix S1: Fig. S18(c)). Salinity, irradiance and wind energy were therefore the drivers in our physics-based models. We note that nutrients, especially N and Si, were also dependent upon freshwater inflow, which explained their high correlation (Appendix S1: Fig. S7).

Most plankton genera showed a strong seasonal response, except for CRY, GUI at Buoy 7 and SKE. The CRY group differed from other algae in its phenology by exhibiting much weaker (if any) periodicity. GUI and SKE may have a stronger long-term (supra-annual) component than other planktonic species (Supplementary S6: Fig. S2(c) and Fig. S2(j)). Finally, shorter cycles (6 months) could be noticed for AST, GUI and NIT; these cycles indicate two blooms per year.

### Nutrient limitations

In a coastal lagoon with continental inputs such as AB, the odds that concentrations in nutrients would be high enough for plankton responses to saturate with regards to nutrient concentration are high. However, using half-saturation values from the literature, we could not show nutrients to be saturating for all genera at both sites (Table 2). Si seemed to be less limiting than N for Arcachon Bay (despite its general ecological importance for diatoms). The high correlation between Si and N, that was more limiting, led us to select N rather than Si as an environmental driver. P seemed to be lacking in comparison with other nutrients, and was therefore considered potentially limiting as well.

### Single-species autoregressive models

When taking all abiotic variables into account, the best model - in the sense of lowest AICc for most planktonic groups used raw values of nutrient concentrations, as opposed to saturating functions of concentrations or nutrient ratios. This “absolute concentrations” model was consistent across genera: 7 genera out of 12 were within *±*2 of the lowest AICc value across all possible model types (including seasonality or not, ratios of nutrient concentrations, saturating functions of nutrient concentrations, or absolute values of nutrient concentrations). The best model also included an explicit seasonal component (Table 3).

**Table 3:** AICc for models explaining plankton dynamics in Arcachon Bay using physical variables (irradiance, wind energy, salinity) and nutrients (nitrogen and phosphorus), with and without a dedicated season variable. Nitrogen values are either used directly, or clipped with a saturating function. Nutrients were also input as ratios of P/N and Si/N. AICc values within 2 units of the minimum AICc are shown in bold letters for each group. Composition of planktonic groups is described in Table 1

|  | No season<br>No saturation | Season<br>No saturation | No season<br>Saturation | Season<br>Saturation | No season<br>Ratio | Season<br>Ratio |
| --- | --- | --- | --- | --- | --- | --- |
| AST | 1430 | 1431 | **1426** | **1427** | 1431 | 1432 |
| CHA | 1708 | **1703** | 1717 | 1706 | 1720 | 1710 |
| CRY | 1187 | **1185** | **1186** | **1185** | **1185** | **1184** |
| EUG | 1005 | 1003 | 1002 | 1002 | **996** | **995** |
| GUI | **1268** | 1272 | **1269** | 1273 | 1271 | 1272 |
| GYM | 1269 | **1260** | 1269 | **1260** | 1266 | 1264 |
| LEP | 1115 | **1109** | 1113 | **1111** | 1113 | 1112 |
| NIT | 1564 | **1545** | 1557 | **1546** | 1561 | **1546** |
| PRP | 1128 | 1129 | 1129 | 1130 | **1121** | **1121** |
| PSE | 1571 | **1566** | 1569 | 1569 | 1569 | 1569 |
| RHI | **894** | **893** | **894** | **894** | **893** | **893** |
| SKE | **804** | 811 | **806** | **806** | 808 | 810 |

To decipher which of the abovementioned factors were most important to plankton dynamics, models with subsets of the environmental data were considered. Four models were evaluated (see Methods for details): the full model (physical drivers + nutrients), a model with physical variables only, a model with nutrients only (Fig. 3). The physics-only model was best - including irradiance, salinity, and wind energy - followed by the full model and finally the nutrient-only model. For the two abundant and widely blooming genera (AST, CHA), the full and physics-only models were nearly equivalent in terms of ∆ AICc but clearly better than the nutrient-only model.

**Figure 3:**
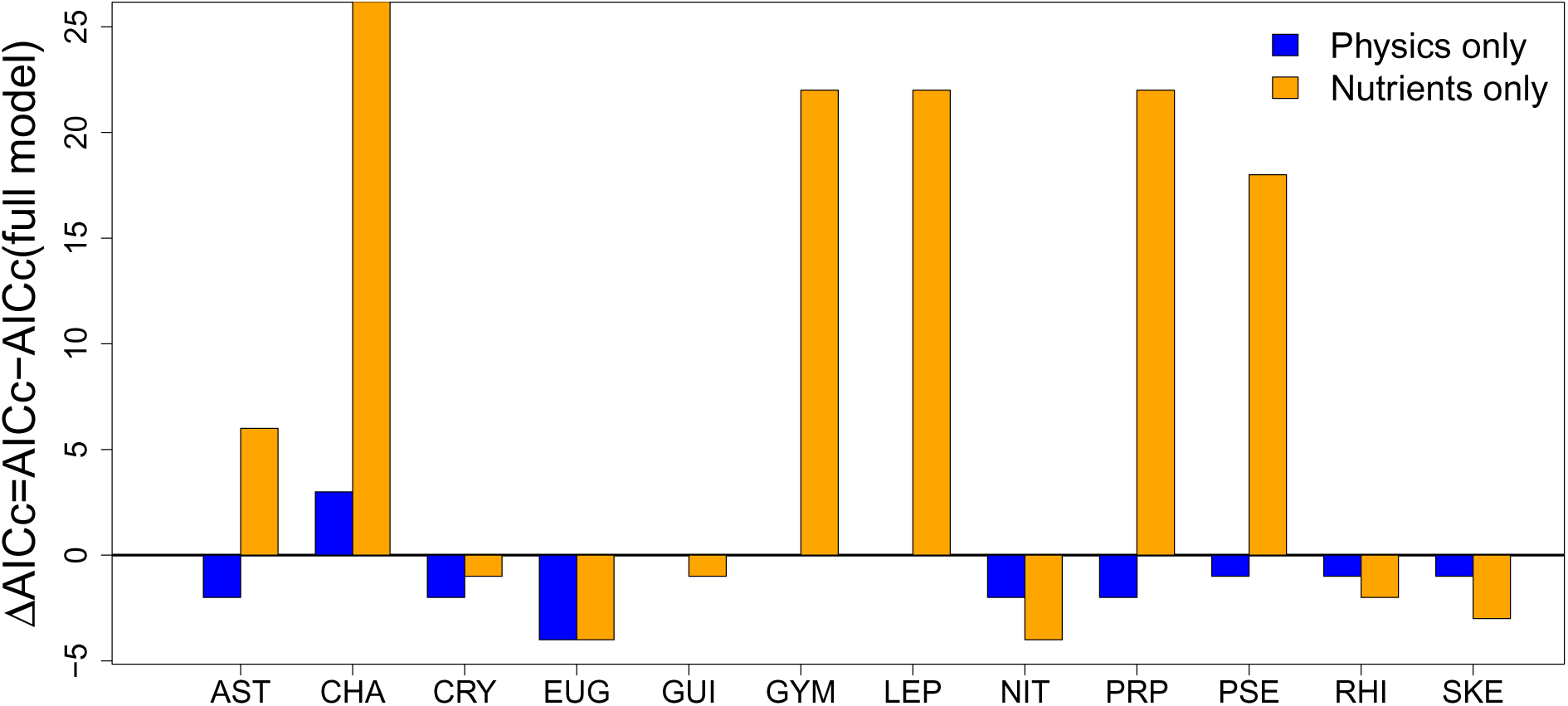
Difference between the AICc of a full model, and the AICc of physics-only (salinity, irradiance, wind) and nutrients-only (nitrogen and phosphorus) models. All models use raw nutrient values and an explicit seasonal component. When ∆AICc is negative, the model performs better than the full model to describes the dynamics of each planktonic groups. Physics-only ∆AICc cannot be seen for GYM and LEP as they are 0. ΔAICc for nutrients-only models were above 40 for CHA. Composition of planktonic groups on the x-axis is described in Table 1.

Even though the physics-only model did not account for all the variability in population growth rates (*R*^2^ ranged between 0.12 and 0.27) and density-dependence was, in general, more influential to explain the overall dynamics than any other environmental factors, some environmental effects such as those of salinity and wind appeared consistent and common to all planktonic groups (Fig. 4). Seasonality had a strong impact on growth rates - the sign of which depends on the life history of the different genera. Residuals from cumulated irradiance had a positive effect, except for AST which is known to be less sensitive to irradiance at Arcachon's latitude (early bloomer), and for PSE and RHI, whose growth rates were driven mostly by variations in wind energy and salinity. Except for a negligible effect on CRY and NIT, wind energy had strong negative impacts on population growth (Fig. 4).

**Figure 4:**
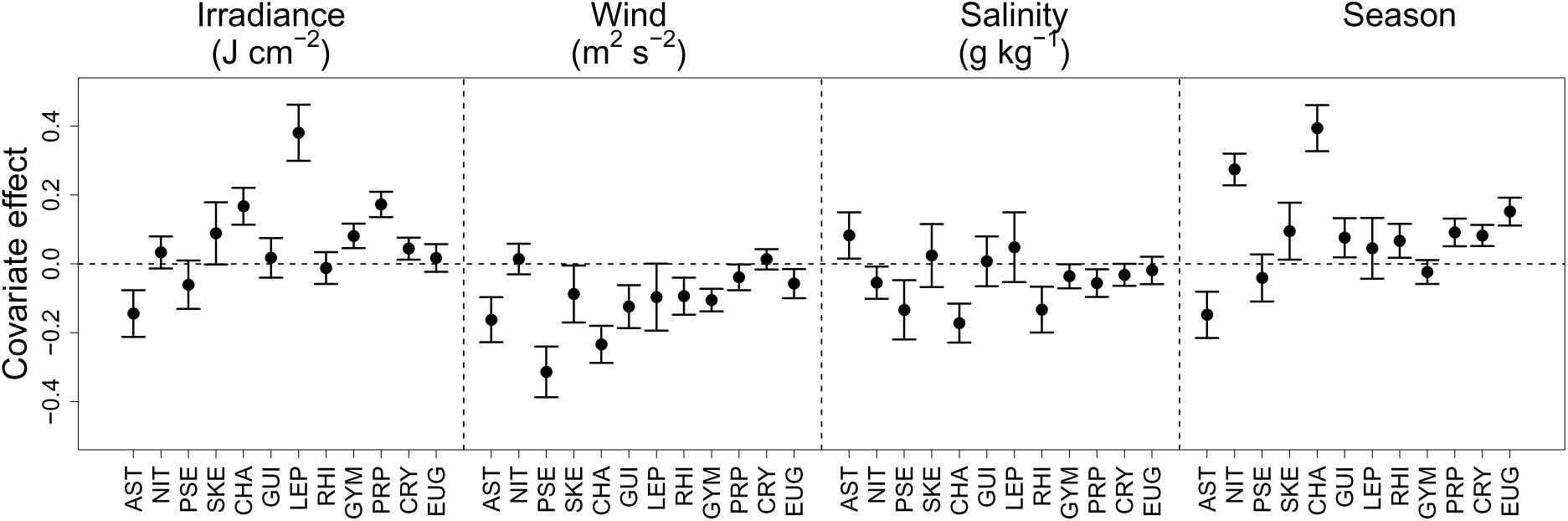
Mean values and standard errors of coefficients from linear models of plankton growth rates against irradiance, wind energy, salinity and a dedicated season component. Composition of planktonic groups on the x-axis is described in Table 1.

The physics-only model was therefore used for MAR analyses in the following.

### MAR(1) models

Model selection with AICc revealed that both the null model (no interaction) and the matrix enabling interactions within centric and pennate diatoms on the one hand, and dinoflagellates on the other hand (eq. 4), were the best models to describe data at Teychan and Buoy 7 sites (Table 4). This was also confirmed by a more detailed analysis of interactions between the main pennate diatom AST and the main centric diatom CHA (see below). **B** and **C** matrices are presented for Teychan and Buoy 7 in Fig. 5 for the model differentiating pennate and centric diatoms. Results based on other interaction matrices can be found in Appendix S2: Section S2.4. In order to make interaction coefficients comparable, the identity matrix was subtracted to all estimates of **B** (see Material and methods: MAR(1) models). Model fit was checked by examination of residuals (Appendix S2: section S2.5) and comparisons of simulated data to real data (Fig. 6).

**Figure 5:**
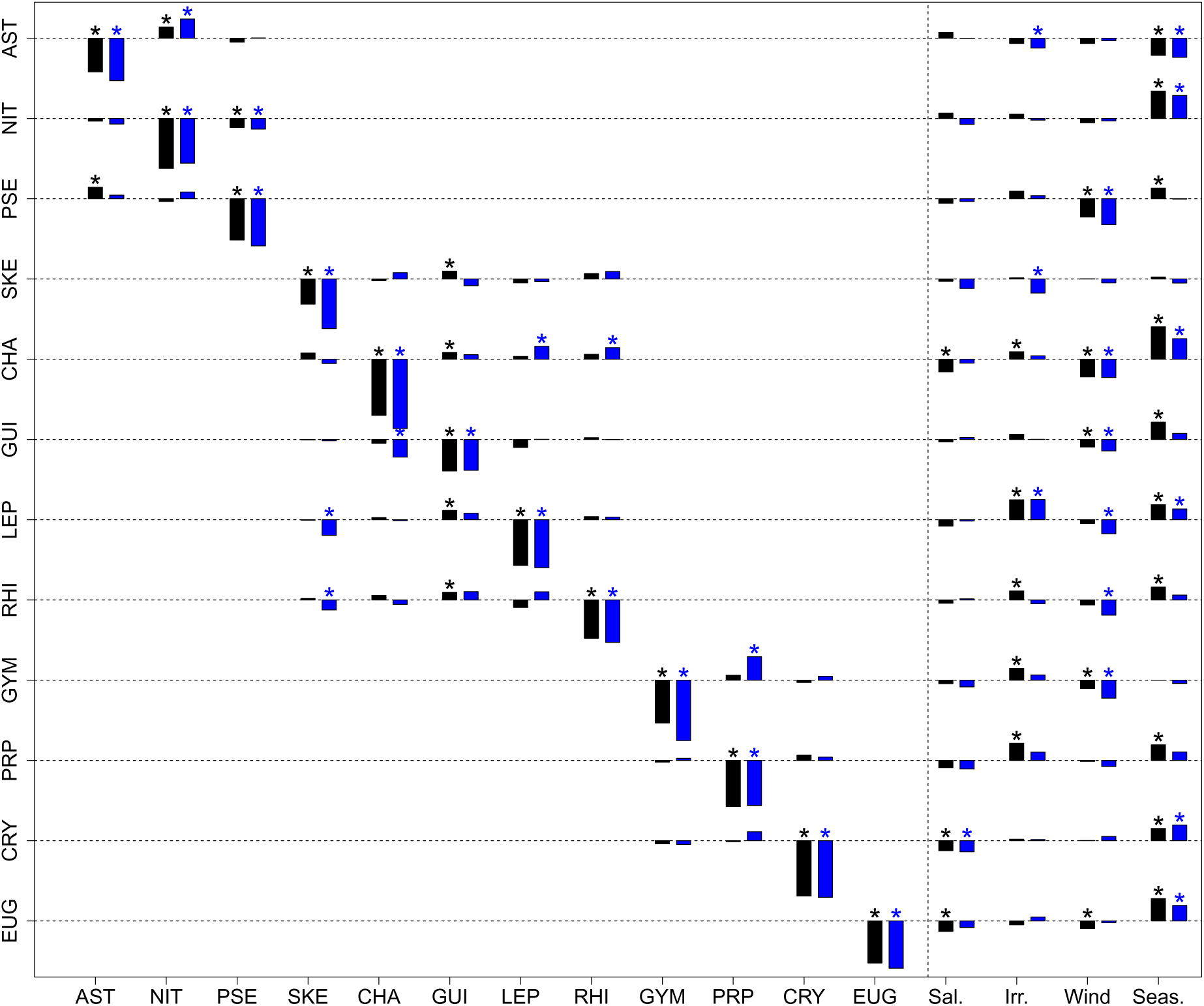
Model coefficients for both variates and covariates (salinity, irradiance, wind energy and season) at Teychan (black) and Buoy 7 (blue), using an interaction matrix with no interaction between pennate and centric diatoms. Significance of coefficients was determined by bootstrapping and P<0.05 and is marked by asterisks (*). The figure should be read as element *i* having effect *e_ji_* on plankton group *j*. The identity matrix was subtracted to the interaction matrix in order to make effects on growth rates comparable. Composition of planktonic groups is described in Table 1.

**Figure 6:**
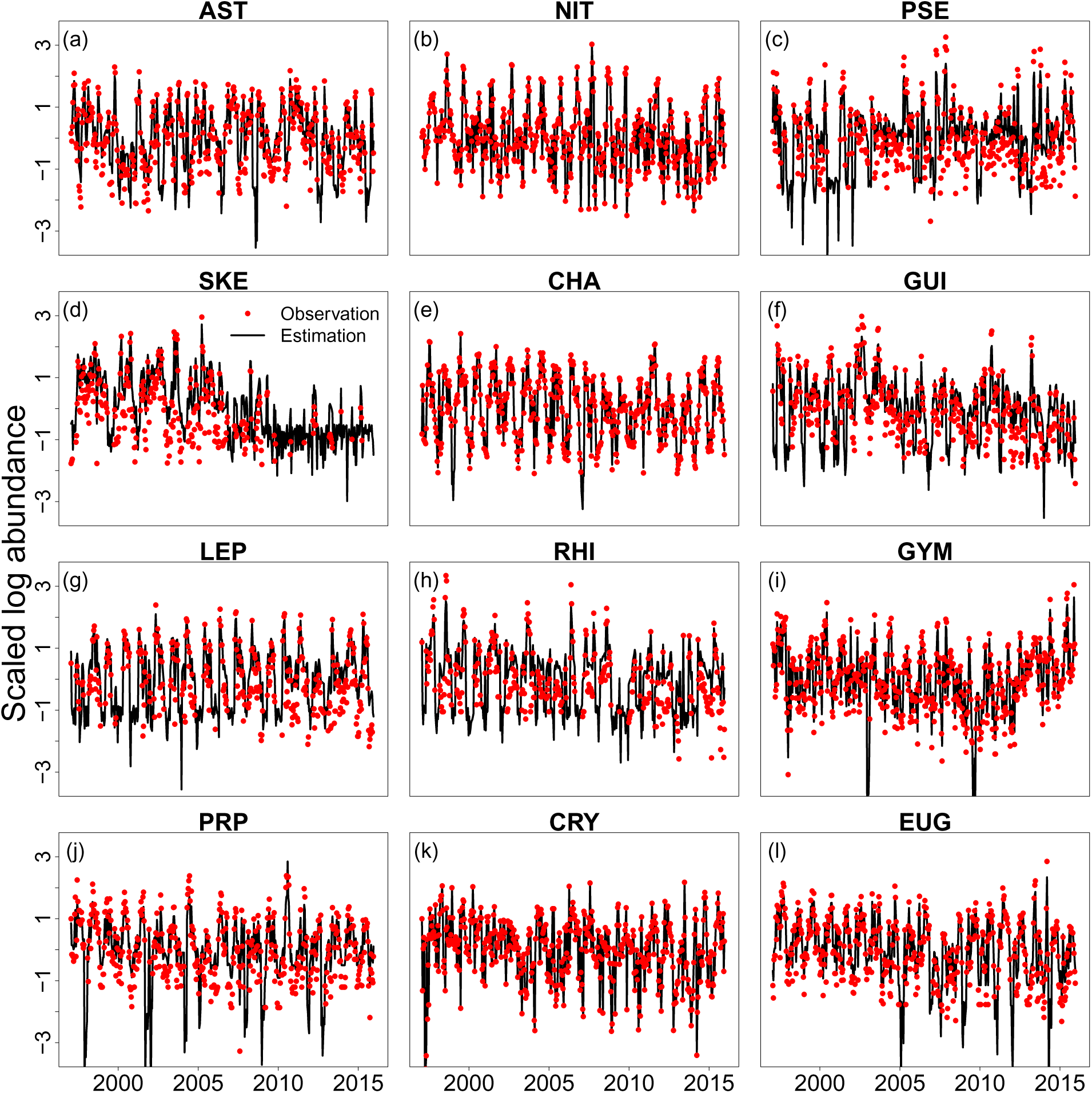
Smoothed states obtained with the Kalman filter used in MARSS package (black line) compared to observations (red dots) for Teychan site, using an interaction matrix with no interaction between pennate and centric diatoms. Composition of planktonic groups is described in Table 1.

**Table 4:** Comparison of model selection criteria for different interaction matrices at Teychan and Buoy 7 sites (see eq. (4) and Appendix S2: Tables S5, S6, S7 for matrix definition)

|  | Null | Unconstrained | Pennate vs. Centric | Diat vs. Dino | Inter-phylum |
| --- | --- | --- | --- | --- | --- |
| Teychan AICc | 11970 | 12039 | **11964** | 11971 | 12025 |
| Teychan BIC | **12271** | 12882 | 12398 | 12533 | 12631 |
| Teychan $R^2_{\text{pred}}$ | 0.52 | 0.55 | 0.50 | 0.50 | 0.54 |
| Buoy 7 AICc | 8924 | 8980 | **8889** | 8920 | 8972 |
| Buoy 7 BIC | **9196** | 9736 | 9280 | 9422 | 9517 |

We found generally that intragroup interactions were at least twice higher than intergroup interactions (often 4 times higher). The interaction matrix structure was found to be similar between sites. Among diatoms, the strongest intragroup regulation was observed for CHA, one of the most abundant genus in AB. AST, the most abundant genus, had an average intragroup interaction coefficient when compared to other planktonic groups. Intergroup coefficients were mostly low and positive when significantly different from zero (Fig. 5). 11 intragroup coefficients were positive at both Teychan and Buoy 7 sites, while only 5 of them were negative and consistent between sites. However, low positive and negative coefficient values might be artefacts, as the model with diagonal **B** was equally supported (Table 4). Covariates played similar roles here than in single-genus models: salinity and wind energy had mostly negative effects while cumulated irradiance had positive effects for the majority of planktonic genera. Our simulations of three scenarios (Appendix S2: Section S2.1) followed by MAR model estimation revealed that while sign and rank of covariate effects were correctly estimated in all scenarios (there could be a slight underestimation bias, c. 10%).

### Bivariate phase-dependent (TAR) models

In this section, we report the results of piecewise autoregressive log-linear (TAR) models, with the phases defined either through the species densities or the general environmental conditions for Teychan (Table 5). Similar results for Buoy 7 can be found in Appendix S5: SectionS1.2.

**Table 5:** Density- and environment-based TAR model coefficients for a linear model relating growth to log abundance of planktonic groups for *Asterionellopsis* (AST) and *Chaetoceros* (CHA) at Teychan, according to Eq. 5, with cumulated irradiance as the main environmental driver. Significant coefficients at the 5% threshold are indicated by *.

| Model | Phase |  | Coefficients |  |  |  |
| --- | --- | --- | --- | --- | --- | --- |
| | | | a | AST | CHA | Irradiance ($\times 10^{-5}$) |
| Density-based | Bloom | AST | 6.38* | -0.60* | 0.05 | -2.98 |
|  |  | CHA | 3.86 | 0.04 | -0.53* | 4.22* |
|  | No bloom | AST | 6.40* | -0.52* | -0.06 | -2.72* |
|  |  | CHA | 4.54* | -0.02 | -0.56* | 5.53* |
| Environment-based | Bloom | AST | 6.47* | -0.53* | -0.05 | -2.94* |
|  |  | CHA | 3.21* | 0.01 | -0.47* | 5.75* |
|  | No bloom | AST | 5.48* | -0.48* | -0.01 | -2.81 |
|  |  | CHA | 5.33* | 0.01 | -0.65* | 4.50* |

The intergroup coefficients are at least two times lower than the intragroup density-dependence (and usually an order of magnitude lower) and statistically non-significant. This was true in both bloom (high densities, or favorable environment) and non-bloom (low densities, or unfavorable environment) conditions at both sites. We therefore conclude that there are very few or no interactions between AST and CHA, even when considering more complex nonlinear models.

## Discussion

Using time series data from both biotic and abiotic components of the ecosystem, we have applied univariate and multivariate autoregressive models to pinpoint the factors driving the joint dynamics of a plankton community in a coastal ecosystem. This plankton community was dominated by several genera of pennate and centric diatoms as well as dinoflagellates (grouped at the genus level for most). Because most of these groups overlap in their resource requirements (Reynolds 2006), we expected some degree of competition but relatively different, potentially nonlinear responses to environmental variables. This would have suggested a mechanism of coexistence through temporal variation of the environment (Chesson 2000), despite some degree of competition between genera. Our results painted in contrast a rather different picture, with no or very weak competition.

Some MAR studies have constrained the signs of the interactions between groups, assuming competitive interactions at the same trophic level (e.g., Ives et al. 2003; Huber and Gaedke 2006; Hampton et al. 2008). We chose not to do so, as it was not clear to us that the resulting net effects of all possible interactions (including those mediated by hidden players such as predators or parasites) are necessarily negative (see also Stone and Roberts 1991). In contrast, we relied on using model selection criteria to contrast the likelihoods of various scenarios (including no interactions between groups, interactions between closely related groups only, interactions between all the plankton groups).

We did not find intergroup competition between diatoms or dinoflagellates. Although one of our best models included interactions between genera (including competition within pennate and centric diatom groups), interaction coefficients that were significantly different from zero and consistent for both sites were usually positive. As the null model without interactions was also selected, we conclude that the odds of no, or weak, interactions between groups are high. Hence, the most likely explanation for our results is that plankton groups currently do not compete to a great extent, even when some groups concurrently reach densities high enough to be considered blooming.

The responses to environmental variables were relatively similar among taxa, but there were some phenological differences. The most abundant group in AB, AST (see Table 1) was a good example of such differences as it almost always bloomed first and necessitated lower irradiance than all other groups. In AB, this group consists almost entirely of *Asterionellopsis glacialis*, a species adapted to cold environments (Kaczmarska et al. 2014). The effects of nutrients were not selected in autoregressive models, even though we considered absolute concentrations, saturating functions of these concentrations, and ratios of nutrient concentrations. There was, however, a possible impact of nitrogen but it was neither high enough nor consistent enough across groups to be considered a key driver, compared to other abiotic factors like irradiance or wind.

Our models relied on the assumption of log-linearity in population growth rates (PGRs), which amounts to assume a multiplicative, power-law equation for the untransformed densities. Both PGR-log(density) curves (Appendix S2: Fig. S1) and examination of residuals (Appendix S2: Section S2.5) did not reject this hypothesis. Because examination of residuals of time series models can sometimes be ambiguous with noisy data, to be sure that alternative modeling choices could not alter our results, we fitted supplementary nonlinear competition models on the two main genera corresponding to pennate and centric diatoms (AST and CHA respectively; see Methods and Appendix S5 for the models definitions). These included Ricker-based Lotka-Volterra (LV) competition (Ives et al. 1999b) (Appendix S5: Section S2) and two phase-dependent models (i.e., piecewise, Stenseth et al. 1998a; Barraquand et al. 2014; Stenseth et al. 2015). The phase-dependent models mimicked relative nonlinearity of competition, by allowing phase to depend on density values, and the storage effect mechanism (Chesson 2000), by allowing the phase to depend on environmental conditions (see Appendix S5 for the definition of environmental conditions). The Ricker-LV model fitted the data to some extent but did not produce realistic dynamics (Appendix S5: Section S1.2) compared to MAR(1) models. The phase-dependent models produced very similar results to the basic MAR(1) log-linear model: no significant interactions were found in any phase. Hence, these models with increased nonlinearity did not bring any additional information. We are therefore very confident that (1) the log-linear assumption is appropriate for our data and (2) there is an absence of interactions between the two most abundant genera (AST and CHA), so that our MAR(1) model including phylogenetic information in the interaction matrix (with weak interactions within pennate and centric diatom groups, as well as dinoflagellates) is quite robust.

We now discuss the likely reasons behind this absence of intergroup competition, as well as the effects of physical environmental drivers on phytoplankton population dynamics.

### Phytoplankton responses to environmental variation

There is a strong influence of the season on phytoplankton growth processes. We have taken into account this variation through an explicit seasonality variable to avoid collinear regressors in our models. Seasonality is explicitely considered in about half of the MAR studies focusing on planktonic communities (Ives et al. 1999a; Klug and Cottingham 2001; Beisner et al. 2003; Hampton et al. 2006; Hampton and Schindler 2006; Hampton et al. 2008; Griffiths et al. 2015; Gsell et al. 2016). It is usually assessed by categorical or continuous variables such as the month, week or day number, sometimes taking into account non-linearities through squared values (Ives et al. 1999a; Hampton and Schindler 2006; Griffiths et al. 2015). More complex analyses have used spline functions to describe the seasonal signal in each covariate (Feng et al. 2014); our choice of a trigonometric function allowed to be more parsimonious. The season variable aggregates, of course, the effect of several other variables that have seasonal variation. Therefore, all the other variables, that are deseasonalized, have to be interpreted as deviations from the overall seasonal pattern.

With these limitations in mind, it is remarkable that both wind energy and salinity had mostly negative effects, meaning that more wind or more salinity than expected at a particular time of the year has detrimental effects on planktonic population growth. Why wind has a consistently negative effect is not entirely clear. Wind has different effects at different spatial scales. At very large spatial scales (>100 km^2^), wind can have disruptive effects on oceanic currents and change the stratification outside coastal areas. In turn, a change in large-scale currents can create favorable conditions for blooms, including in the Bay of Biscay where AB is located (Díaz et al. 2013). At slightly smaller scales (m^2^ to km^2^), wind is generally expected to create large-scale turbulence, which is then cascaded to small-scale eddies and dissipated as heat at the micrometer scale. Despite the development of multiple modeling studies (Huisman et al. 1999a; Guasto et al. 2012; Nguyen and Fauci 2014), field observations and experiments in the laboratory are not conclusive on the effect of small-scale turbulence (providing mixing and heat), generated by larger-scale phenomena (Peters and Marrasé 2000). Positive effects of microscale turbulence on planktonic growth include lowered sinking rate for diatoms (Reynolds 2006) enhanced by increased aggregation and improved nutrient assimilation (Margalef 1978; Thornton 2002). However, other studies have shown that an increase in flow velocity result in decreased growth rates and increased mortality of phytoplankton (Li et al. 2013; Garrison and Tang 2014), especially for dinoflagellates (Peters and Marrasé 2000; Llaveria et al. 2009). Moreover, an increase in turbulence may lead to resuspension of sediments and therefore more turbid waters and lower light availability. The buffering of incoming solar energy could explain the observed negative effect of wind energy on plankton growth (Gervais et al. 1997; Glé et al. 2007).

Salinity is another dominant factor in other coastal areas, that can prove at times even more influential than nutrient loads (Irwin et al. 2012; Gasiūnaitė et al. 2005). Scheef et al. (2013) called for the use of salinity as a discriminating factor for estuarine environments in MAR analyses. Some diatoms have indeed lower growth rates when salinity increases (Balzano et al. 2011), which could explain our mostly negative effects on plankton growth. An alternative hypothesis is that salinity is inversely related to freshwater inflow and therefore nutrient loads. As we found few direct effects of nutrients, it seems more logical to assume that salinity might have a negative effect for other reasons than its inverse relationship to nutrient load. Salinity is therefore most likely to influence phytoplankton growth directly through physiological adaptations (e.g., tolerance range due to osmotic processes, Kirst 1990) or indirectly through concentration effects other than nutrients (e.g., a higher freshwater inflow can dilute chemicals harmful to algae).

### No (or weak) interspecific competition

The absence of competition in our system was surprising, because we expected to find results similar to Descamps-Julien and Gonzalez (2005), i.e., a coexistence in spite of marked competition, mediated by differential responses to environmental variation. We note that our detailed comparison of nonlinear community models for AST versus CHA is not unlike their comparison of *Fragilaria* (pennate diatom) vs *Cyclotella* (centric diatom), except ours was based on field-based rather than based on experimental data. Although the absence of present competition between groups could be expected because niche differentiation has occurred in the past in many communities, it is not quite obvious what these niches may be here.

All our plankton groups coexist in a bay where spatial variation at large (e.g., meter to kilometer scale) is low, the resource (nutrient) requirements are mostly met and the responses to these resources are both weak and similar. Our observational setting contrasts widely with the classic experiments of Titman (1976); Tilman et al. (1982) that showed competition for resources in a context with well-defined niches related to nutrients. It is possible that under the differing circumstances considered here, entirely new drivers, e.g., predators or parasites take over the regulation of phytoplankton growth. We discuss this possibility further below.

Among the few intergroup interactions that were consistent across sites, we observed more positive than negative interactions. This can be due to hidden players that generate positive interactions (see below). Although we reckon that there is a possibility for positive interactions to emerge if the seasonal variable does not fully correct for shared seasonal trends, positive interactions can also be genuine ecological phenomena (de Ruiter and Gaedke 2017). A number of plankton MAR studies have explicitely forbidden positive interactions between phytoplankton groups from their models (e.g., Ives et al. 2003; Huber and Gaedke 2006; Hampton et al. 2008), which in turn suggests that positive interactions could be more common than what previous multivariate time series analyses concluded.

### What about neutral dynamics?

Neutral dynamics sensu Hubbell (2001), i.e., ecological drift in a zero-sum game, is likely not a main force here because of the fluctuations in numbers across 6 orders of magnitude, as we mentioned in our introduction. However, there are other models of neutral community dynamics that may apply to our system, such as the one proposed by Loreau and de Mazancourt (2013). It is a variant of the discrete-time Lotka-Volterra model (Ives et al. 1999b) with demographic and environmental stochasticity. In the neutral case, the species compete together by an equal amount, yet vary synchronously under the influence of the environment. Stabilizing niche differences are progressively introduced through a parameter that weights the importance of interspecific/intergoup interactions. Our results may be relevant in two ways for such theoretical community dynamics studies.

First, in our study, not only species vary in partial synchrony under a joint environment, but intragroup interactions also dwarf intergroup interactions (i.e., close to zero in the second model of Loreau and de Mazancourt 2013). In this case, the Loreau and de Mazancourt (2013) model is not neutral anymore and it contains stabilizing mechanisms for coexistence (sensu Chesson 2000). In our case, the reasons for such stabilizing niche differences are yet to be discovered. Whether other studies using long time series, in plankton or other similar systems find results close to ours, or manage to get near-neutral dynamics, would be of general interest. If coexistence occurs through stabilizing niche differences, we note that observing intergroup interactions that are very small may be in fact logical: the criteria for coexistence in theoretical Lotka-Volterra communities require intergroup competition to be much lower than intraspecific competition in large communities (Barabás et al. 2016).

Mutshinda et al. (2016) recently reported on ecological equivalence - neutrality - between groups using like us a long-term phytoplankton dataset (the L4 dataset in the English Channel). Ecological drift was modeled by assuming that the proportion of the functional group biomass attributed to each species was on average the same proportion realized in the previous week. Though we command the extensive hierarchical modeling of Mutshinda et al. (2016), it is not entirely clear to us to which neutrality concept this proportion modeling refers to, and we therefore prefer to interpret our results by comparison to the framework of Loreau and de Mazancourt (2013), where the assumptions on competition are clearly defined and the model structure match the models that we fitted. Better understanding the relationship between “neutrality” sensu Loreau and de Mazancourt (2013) and Mutshinda et al. (2016) might help understand if the L4 phytoplankton community is ‘neutral’ within phyla (diatoms vs dinoflagellates, see Mutshinda et al. 2016 for details) in the sense of Loreau and de Mazancourt (2013) as well. If this was the case, the L4 phytoplankton community would have widely different community dynamics than AB where stabilizing mechanisms dominate.

Our second insight is that we found a clear log-linearity of the relationship between population growth rates and population densities (Appendix S2: Fig. S1 and Appendix S2: Section S2.5), i.e.

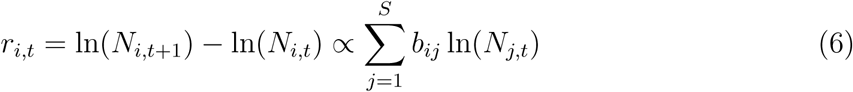

This is the MAR formulation with Gompertz density-dependence, rather than Ricker density-dependence (used for discrete-time equivalents to the Lotka-Volterra model) as in Loreau and de Mazancourt (2013)

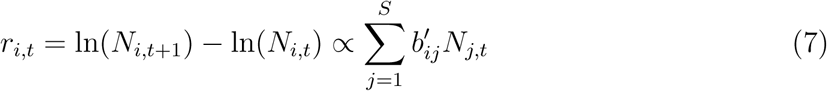

where 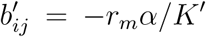 when *i* ≠ *j* using Loreau and de Mazancourt (2013)'s notations. We therefore suggest to theoreticians and empiricists alike to use, when investigating neutral or potentially Lotka-Volterra dynamics in highly variable communities, the Gompertz formulation in addition to the Ricker formulation (Ives et al. 1999b).

### Comparison to biotic drivers in similar ecosystems, and taxonomic issues

We found no interaction between diatoms and our mixotrophic/heterotrophic dinoflagellates, as did for example Gsell et al. 2016. Such interactions between classes are highly variable in other studies: Scheef et al. (2012) only found a positive effect of dinoflagellates on diatoms while Griffiths et al. (2015) estimated a negative effect of autotrophic and mixotrophic dinoflagellates on diatoms in their coastal site and a positive effect of diatoms on heterotrophic dinoflagellates in their offshore site (c. 40km away). In addition, in our study, cryptophytes do not affect and are not affected by other plankton groups at both sites, which surprised us; we expected this abundant group with known interactions to other group to exhibit some significant interactions. In another plankton study, Klug et al. (2000) found a negative effect of cryptophytes on dinoflagellates in one location and no effect in another site (Klug and Cottingham 2001). Basic ecological information suggest that dinoflagellates can have a negative effects on cryptophytes population dynamics (through predation, Moeller et al. 2016), but these seem to be quite difficult to recover.

The above summary of previous plankton MAR studies suggest that patterns of interactions between broad plankton groups at the class or phylum level (diatoms / dinoflagellates / cryptophytes …) are highly inconsistent between places and periods of study. These idiosyncratic results might be due - among other factors - to the aggregation of heterogenous genera and species with different traits and dynamics. For this reason, Griffiths et al. (2015) suggested that improving taxonomic resolution, as we did, would help to estimate more precisely competitive or predatory interactions between plankton groups. However, we did not estimate stronger interactions despite having quite long time series, which suggests that interspecific competition at least may in fact not exist in many (phyto)plankton datasets.

Are there other datasets corroborating our findings? Our inference of interactions using MAR models on field data, at a fine taxonomic scale, can to our knowledge only be compared to the detailed freshwater study of Huber and Gaedke (2006). Distinguishing 4 diatom species, 2 cryptophyte groups and 1 dinoflagellate species, they could infer a negative effect of a dinoflagellate on one species of diatom. However, a different species of the same diatom genus was not affected by the dinoflagellate. Species-specific interactions could also be found between diatoms and cryptophytes. One of our main results was the absence of interactions between pennate and centric diatoms. Interspecific interactions within diatoms were only found between one centric species and one pennate species in Huber and Gaedke (2006), but were not a common pattern; thus their results tend to concur with ours. Although we reckon that our aggregation to the genus level - still more precise than most similar MAR studies - could remove some of the temporal variability at the species level (see also Vasseur and Gaedke 2007).

New techniques to structure interaction matrices applied to metabarcoding data may hold promise to infer interactions at the species level (Ovaskainen et al. 2017) in the future, although currently metabarcoding presents its unique set of challenges, such as data compositionality (Cao et al. 2017). In any case, we recommend that future studies use simulations mimicking study design as we (Appendix S2: Section S2.1) and Ovaskainen et al. (2017) did - simulations are essential to check that statistical models are able to infer interactions for a given observational study design in a range of scenarios.

### Hidden (biotic) drivers

Effects of phytoplankton natural enemies can be quite important (Klug et al. 2000; Ives et al. 2003; Huber and Gaedke 2006; Gsell et al. 2016) but were unknown for AB. There is only one study on the effect of viruses in AB, which shows no viral control of plankton but a high potential for viral lysis during blooms (Ory et al. 2010). Zooplankton is another potential compartment whose populations show a strong spatial and temporal heterogeneity both in biomasses and species diversity from oceanic waters to coastal waters, which could explain some of the differences in coefficient estimates between our two study sites (Castel and Courties 1982; Tortajada et al. 2012). Predators can even contribute to apparent facilitation between species, therefore explaining the positive interactions found in our study (Abrams et al. 1998; Barraquand et al. 2015; de Ruiter and Gaedke 2017). However, in a well-mixed environment, sharing predators can also be like sharing resources, and the species that resist better predation can end up excluding the others, depending on predator numerical responses (Holt 1977; Abrams et al. 1998). Thus natural enemies do not necessarily have diversity-enhancing effects that explain the coexistence observed here. We mentioned parasitism, predation and competition with other algae as potential factors that should be taken into account (Huber and Gaedke 2006), but other variables may emerge as potential drivers of plankton dynamics, from toxic compounds to hydrodynamic features.

### Comparison to abiotic drivers in similar ecosystems

In Arcachon Bay, Glé et al. (2007) focused on the drivers of the winter bloom with a 5-year data set and showed the importance of irradiance for planktonic blooms without being able to conclude on the role of the nutrient cycle throughout the year based on their observations. A high-frequency, detailed sampling program was then launched for two years but its conclusions are difficult to generalize, due to the occurrence of a heat wave in the study period (Glé et al. 2008). With a longer time series (1993-2010), David et al. (2012) classified planktonic groups with a functional approach and indicated that climate indices (NAO and AMO) were more informative than nutrients when observing planktonic diversity patterns along the French coast. Likewise, our study showed no dominant effect of nutrients in AB, which is similar to what Griffiths et al. (2015) found for diatoms and cryptophytes in the Baltic sea. A potential competition for nutrients between phytoplankton, seagrass and macro-algae was suggested for AB based on mechanistic modeling (Plus et al. 2015), but as we did not find nutrient limitation in our analyses, this hypothesis currently seems unlikely. In contrast to small-scale experiments, analyses of phytoplankton time series based on long-term observational data rarely show a consistent effect of nutrients on diatoms (no effect in Klug and Cottingham 2001; Hampton and Schindler 2006, contrasted effects in the three time periods considered by Gsell et al. 2016). These inconsistent nutrient effects may occur because effects of nutrients on phytoplankton growth interact with other important drivers of phytoplankton community dynamics in the field (as opposed to the more controlled environments in which experiments are usually performed).

### Towards a more mechanistic study of phytoplankton community dynamics

In land plants, coexistence can be high-dimensional and depend on a great variety of traits, which are themselves not directly related to abiotic resources such as nitrogen or phosphorus (Kraft et al. 2015). This is also quite likely to happen for phytoplankton. The phytoplankton groups that we studied, diatoms in particular, often live in colonies of differing shapes and sizes: these factors are likely to impact community dynamics and coexistence. The colonial nature and shapes of diatoms are known to interact with patterns of very-small scale turbulence (Margalef 1978; Reynolds 2006), and microscale hydrodynamics does affect plankton growth and coexistence (Huisman et al. 1999b). Much in the same way that small-scale spatial structure can be pivotal to plant coexistence (e.g., through vertical shading, Onoda et al. 2014), the interplay between microscale hydrodynamics and phytoplankton structure (cell shape, cell size, coloniality) is likely to be important for phytoplankton coexistence. Current models have started to investigate the joint effects of cell size and turbulence on persistence (Portalier et al. 2016), but much remains to be done with respect to cell shape and coloniality (see Nguyen and Fauci 2014 for a hydrodynamics study).

This study highlighted that stabilizing niche differences, that make intragroup competition much stronger than intergroup competition, are likely at work in our coastal phytoplankton community. This contrasts widely with other types of models invoking temporal fluctuations in the environment (Descamps-Julien and Gonzalez 2005; Li and Chesson 2016) that counteract effects of competition for resources. However, this rejection of “paradox of the plankton models” does not imply that all interactions between species are unimportant to species coexistence. Other features of phytoplankton life histories may need to be explored to better comprehend the full effect of interactions on plankton interlinked population dynamics, e.g., sexual reproduction in phytoplankton ensures a rather rich demography (D'Alelio et al. 2010). Having several life-stages increases the possibility for interactions to affect community dynamics without these interactions being detectable using aggregated biomass or counts at the species or genus level (Oken and Essington 2015) - one of the limitations of count-based studies like ours.

To sum up, both demography and microscale hydrodynamics, together with natural enemies, might be needed to understand coexistence in phytoplankton. Although the sessile nature of land plants and the stability of forests contrasts with the extremely motile, dynamic nature of the marine environment, some suggestions for further work may be transferrable from plant to plankton ecology. A likely driver of stabilizing niche differences in tropical forests are Janzen-Connell effects where greater interspecific than intraspecific competition occurs at the seed and seedling stage, through combined effects of spatial structure of the seed rain and natural enemies (Bagchi et al., 2014; Comita et al., 2014). Although the specific mechanisms at work will undoubtedly be different in phytoplankton, the idea that microscale spatial structure, stage structure and natural enemies may all interact to create stabilizing niche differences holds promise. A more demographic, small-scale perspective may help us to move beyond simple “paradox of the plankton” models and better explain the puzzling phytoplanktonic diversity.

## Acknowledgements

This study could not have been performed without the dedicated, long-term data collection of the REPHY programme undertaken by Ifremer, for which we are deeply grateful. We would like to thank in particular Nadine Neaud-Masso, Myriam Rumèbe, Claire Méteigner for phytoplankton counts and Florence d'Amico for hydrological measurements. We also thank Météo France for meteorological data. Grégoire Certain, Florian Hartig and Jarad Mellard provided constructive comments on the manuscript. This study was supported by the French ANR through Labex COTE (ANR-10-LABX-45). FB and CP have equally contributed to this work.

## Footnotes

1 http://envlit.ifremer.fr/infos/rephy_info_toxines

2 http://www.cpc.ncep.noaa.gov/products/precip/Cwlink/pna/nao_index.html accessed on 2016/03/22

3 http://www.esrl.noaa.gov/psd/data/timeseries/AMO/ accessed on 2016/06/30

4 https://github.com/fbarraquand/PhytoplanktonArcachon_MultivariateTimeSeriesAnalysis This release will be made public upon publication.

